# Evaluation of the accuracy of a smartphone-based artificial intelligence system, PlantVillage *Nuru,* in diagnosing of the viral diseases of cassava

**DOI:** 10.1101/2020.01.26.919449

**Authors:** Latifa Mrisho, Neema Mbilinyi, Mathias Ndalahwa, Amanda Ramcharan, Annalyse Kehs, Peter McCloskey, Harun Murithi, David Hughes, James Legg

**Affiliations:** IITA, Dar es Salaam, Tanzania; University of Dar es Salaam, Tanzania; Pennsylvania State University, State College, USA

**Keywords:** Cassava, CMD, CBSD, PlantVillage Nuru

## Abstract

**Premise of the study:** *Nuru* is an artificial intelligence system for diagnosis of plant diseases and pests developed as a public good by PlantVillage (Penn State University), FAO, IITA and CIMMYT. It provides a simple, inexpensive and robust means of conducting in-field diagnosis without requiring internet connection and provides real-time results and advice. The present work evaluates the effectiveness of *Nuru* as an in-field diagnostic tool by comparing the diagnosis capability of *Nuru* to that of cassava experts (researchers trained on cassava pests and diseases), agricultural extension agents and farmers.

**Methods:** The diagnosis capability of *Nuru* and that of the assessed individuals was determined by inspecting cassava plants in-field and by using the cassava symptom recognition assessment tool (CaSRAT) to score images of cassava leaves.

**Results:** Nuru’s accuracy for symptom recognition when using six leaves (74 - 88%, depending on the condition) was similar to that of experts, 1.5-times higher than agricultural extension agents and two-times higher than farmers.

**Discussion:** These findings suggests that Nuru can be an effective tool for in-field diagnosis of cassava diseases and has a potential of being a quick and cost-effective means of disseminating knowledge from researchers to agricultural extension agents and farmers.

## INTRODUCTION

The steady increase in the world population and changes in climate are adding pressure to agriculture as the need to produce more food intensifies. South Asia, Latin America and Sub-Saharan Africa are some of the regions facing the detrimental effects of climate change due to changes in the frequency and severity of drought and floods in addition to increasing temperatures and decreasing soil water (FAO et al 2018). Increase in temperature is also anticipated to result in increased abundance of many insect pests through higher rates of growth and population development as well as their spread and migration (Lin 2011). Great efforts are therefore being put into enhancing the management of emerging pests and diseases along with the production of climate-resilient and disease-resistant crops in these regions in an attempt to reduce the risk of hunger (FAO et al 2018).

Tools and technologies that can be used for early detection and diagnosis of crop diseases and presence of pests are being encouraged to facilitate management of pests and diseases. For this reason, several developing countries in Africa, Asia and Latin America have adopted the use of ICT-based technologies and platforms in form of the internet, call-centres and SMS to disseminate agricultural information to farmers (Qiang et al., 2012). These technologies have shown promising results in reducing the knowledge gap between experts and farmers as they allowed for transfer of basic skills, state-of-the-art technologies and production techniques (Qiang et al., 2012). However, most of these technologies have not managed to persist in some regions mainly as the tools failed to meet some of the socio-demographic needs of farmers. Examples of the socio-demographic and infrastructural hindrances include language, illiteracy, network availability and electricity as well as funds to afford and maintain the technologies (Aker, 2011).

*Nuru*, a mobile-based app for diagnosis and management of plant diseases and pests, is one of the new and cost-effective ICT-based technologies that can deal with some of the challenges encountered by small scale farmers, thereby increasing their knowledge and helping them to improve their crop production. *Nuru* was created from a deep learning object detection model that can determine the presence of diseases and pests in plants based on foliar symptoms (Ramcharan et al., 2017; 2019). Currently *Nuru* is trained to identify the presence of the viral diseases of cassava (CMD and CBSD) and the damage caused by cassava green mites (CGM) as well as the presence of Fall Armyworm (FAW) on maize based on symptoms that can be seen on the leaves. Together, CMD and CBSD are causes of the major economical losses in cassava production (Legg et al. 2015), while FAW has been reported to cause damage in several major crops including maize, sorghum, rice and sugarcane (Abrahams et al. 2017). On-going work is being done to develop models for diagnosis of diseases on other food crops including potatoes and bananas/plantain.

The cassava model was developed using machine learning algorithms that have been trained to identify symptoms of CMD and CBSD as well as damage caused by CGM on leaf images from different varieties of cassava plants grown at field sites in coastal Tanzania (Ramcharan et al. 2019). The cassava model was subsequently deployed as a mobile app in Android smartphones and then tested in fields to account for environmental factors and hence determine the best conditions that will enable the model to perform with high accuracy.

*Nuru* requires users to have an android smartphone with random access memory (RAM) of 2GB or more. These phones can be obtained at a cost of about US$100-US$150, depending on the region, which can be afforded by some farmers and most agricultural extension agents who serve as a direct link to the majority of farmers in sub-Saharan Africa. Newer and yet more affordable and feature-rich phones are constantly becoming available due to the rapid rate of technological advance in mobile phone development. For this reason, the price of phones compatible with *Nuru* is expected to decrease thereby increasing the accessibility of the app.

*Nuru* can be downloaded, free of charge, from the Android PlayStore and once installed, it can operate in the absence of the internet, enabling access to the tool even in remote areas with poor mobile and internet network connectivity. However, this feature is only available for the diagnosis tool and advice on management pest and disease but not for the additional features that connect the user to researchers, experts and farmers. These additional features include a discussion forum and a platform the enable the user to ask questions or share information with experts who may be able to assist in the diagnosis of the condition of the plant.

*Nuru* is programmed in multiple languages including English and Swahili which are spoken in East and Southern Africa, while ongoing efforts are being made to provide access in multiple other languages through a crowd-sourcing translation tool. Voice command functionality is being integrated into the app to give access to users who are not able to read and write.

*Nuru* gives real-time diagnosis and then advice on management of the disease or pest it has detected enabling the user to deal with the disease or pest in their farms early on and hence prevent spread and higher yield losses. During diagnosis, *Nuru* shows the user the symptoms it has recognised, therefore it can also be used as a teaching tool. The teaching capability of *Nuru* has been enhanced by linking the app with the PlantVillage database where the user can access a library of information on different crops relating to agricultural practices, diseases and pests as well as their management techniques.

The work described here examined the effectiveness of *Nuru* as an in-field diagnostic and teaching tool by comparing the diagnosis capacity of *Nuru* to that of experts in cassava pests and diseases as well as agricultural extension agents and cassava farmers.

## MATERIALS AND METHODS

*Nuru* was evaluated by comparing its capability to diagnose cassava diseases to that of experts, researchers, agriculture extension officers and farmers. This required the experts to be highly effective in determining the presence of diseases on cassava based on foliar symptoms. Hence the capability of cassava experts (from the International Institute of Tropical Agriculture (IITA), Dar es Salaam, Tanzania) to identify foliar symptoms of cassava diseases was determined and then compared to the disease recognition capabilities of extension agents, farmers and *Nuru*.

### Determination of the capability of cassava experts to diagnose cassava diseases based on foliar symptoms

The ability of the cassava experts to accurately identify the symptoms of CMD, CBSD and CGM-damage, based on symptoms observed from the leaves, was assessed by comparing visual and molecular diagnosis (through PCR or RT-PCR amplification of the disease-causing viruses). Two researchers, with combined experience of three years of work on cassava, visually diagnosed 75 cassava plants of five different varieties by assessing the presence or absence of symptoms for CMD and CBSD. The experts randomly selected two leaves (one from the top and the other from the bottom part of each plant), scored the leaves based on the observed condition and then collected the leaves which were used to check for the presence of the disease-causing virus by molecular diagnosis (described in supplementary material A). The accuracy of expert diagnosis was determined by calculating the proportion of plants for which expert’s symptom diagnoses matched those of the virus test results.

### Evaluation of in-field accuracy of cassava AI Nuru in diagnosis of cassava diseases based on foliar symptoms

The ability of *Nuru* to identify symptoms of CMD, CBSD and CGM damage was tested in the field using 90 plants that were visually diagnosed by the experts to be asymptomatic and symptomatic for CMD, CBSD and CGM prior to diagnosis by *Nuru*. During diagnosis with *Nuru*, the app was pointed at the leaves and the symptoms that were detected, identified by boxes that popped up on the diagnosis screen, were recorded. The ability of *Nuru* to accurately diagnose the condition of the plant using multiple leaves was determined by comparing the accuracy of diagnosis obtained when *Nuru* was tested using six leaves (three top leaves and three bottom leaves). The comparison was done by determining the likelihood of the app to give a correct diagnosis when only one or multiple leaves were diagnosed (from upper and/or lower parts of the plant).

### Evaluation the accuracy of the cassava AI Nuru using the symptom recognition assessment tool

Evaluation and comparison of the capability of *Nuru* to that of agricultural extension agents and cassava farmers was done using the symptom recognition assessment tool. This tool was developed at IITA and consist of a scoring matrix for assessing the condition of 170 randomly-selected images of asymptomatic and symptomatic cassava leaves based on the symptoms observed on the leaves. The scores were analysed by the assessment tool which compares the scores from the individual being evaluated to the consensus expert score and gives a percent accuracy for symptom recognition. The assessment tool can also identify incorrectly diagnosed images and identify which condition(s) have been misdiagnosed. Detailed information on how the symptom recognition assessment tool was developed can be found in supplementary material B.

The assessment tool was used to compare the ability of *Nuru* and that of researchers, agricultural extension agents and farmers. Each of the groups (researchers, extension agents and farmers) contained 22 individuals, half of which were individuals who had previous experience of working with cassava and other the other half who did not. Each individual was shown 170 images of cassava leaves (at an interval of 15 seconds per image) and was requested to score the images based on the symptom/condition they observed. The range of images included leaves with symptoms of CMD, CBSD and CGM damage as well as symptomless leaves. *Nuru* was also evaluated on the same images by pointing the cassava AI at a laptop screen showing the images from the cassava symptom recognition assement tool. Boxplots, single factor ANOVA and Tukey HDS statistics were used to evaluate and compare the accuracy scores obtained from the researchers, agricultural extension agents, farmers and *Nuru*. The statistical analyses were done on R Studio Version 1.1.456 (RStudio Inc).

### Evaluating the teaching capability of the cassava AI Nuru

The teaching capability of *Nuru* was evaluated by determining the ability of agricultural extension agents to identify symptoms of the cassava diseases prior to and after using *Nuru*. The disease diagnosis ability of the agricultural extension agents (30 individuals) and farmers (50 individuals) was determined using the cassava symptom recognition assessment tool prior to introduction and usage of *Nuru*. After the assessment, all the agricultural extension officers and farmers were trained on diseases and pests of cassava and then introduced to *Nuru*. Afterwards, both the agricultural extension agents and farmers were divided into two groups, 15 individuals for the groups of agricultural extension agents and 25 individuals for the groups of farmers. One group of farmers and agricultural extension agents was given phones with access to *Nuru* and requested to inspect and collect data from 100 healthy and diseased cassava plants using *Nuru*, within two weeks. The remaining groups of agricultural extension agents and farmers were not given access to *Nuru* nor asked to collect data from cassava plants. After the two weeks, the symptom recognition ability of both agricultural extension agents who were requested to inspect plants with *Nuru* and those who were not, were evaluated using the cassava symptom recognition assessment tool and their symptom recognition accuracy scores were compared.

## RESULTS AND DISCUSSION

### Accuracy of cassava experts in diagnosing symptoms of CMD and CBSD

Comparison of the molecular and visual diagnosis showed that the experts in cassava pests and diseases could determine the presence or absence of either CMD and CBSD in a plant with high accuracy (95% for CMD and 81% for CBSD), when only one condition is considered as illustrated in Table 1.

**Table 1.** Comparison of visual and molecular diagnosis of CMD and CBSD on cassava plants by experts working on cassava pests and diseases.

| Condition | Visual | Molecular | Number of plants | Score (%) | Overall score | Overall condition |
| --- | --- | --- | --- | --- | --- | --- |
| CMD | Present | Present | 19 | 25 | 95% | Accurately diagnosed |
|  | Absent | Absent | 52 | 69 |  |  |
|  | Absent | Present | 3 | 4 | 5% | Misdiagnosed |
|  | Present | Absent | 1 | 1 |  |  |
| CBSD | Present | Present | 35 | 47 | 81% | Accurately diagnosed |
|  | Absent | Absent | 26 | 35 |  |  |
|  | Absent | Present | 9 | 12 | 19% | Misdiagnosed |
|  | Present | Absent | 5 | 7 |  |  |

The majority of the misdiagnoses were due to the presence of CBSD-causing viruses (Cassava Brown Streak Ipomoviruses - CBSIs) that produced no symptoms, also known as latent infection. Patterns of CBSD symptom expression (including the absence of symptoms) are known to change over the course of CBSD infection in cassava plants (Nichols, 1950). Furthermore, the expression of foliar symptoms has been reported to vary between leaves on infected plants, cassava variety, growing conditions (temperature, rainfall, and altitude), age of the plant and the virus isolate involved in causing the disease symptoms (Hillocks and Jennings, 2003; Mohammed et al., 2012).

So it is unsurprising that experts were relatively less successful in correctly diagnosing CBSD infection as compared to CMD. Inability to recognize CMD symptoms was also thought to be due to latent symptoms or limitations in the molecular diagnosis technique used as a confirmatory test for the presence of viruses that causes CMD (Cassava Mosaic Begomoviruses - CMGs).

Since some of the varieties that were tested were infected by viruses causing both CMD and CBSD, the ability of the cassava experts to diagnose symptoms of co-infections of CMD and CBSD was also evaluated. The accuracy of the cassava experts to diagnose healthy plants (asymptomatic) and symptoms of CBSD in CBSD-infected plants was higher (82% and 90%) than that for CMD-infected and CMD-CBSD co-infected plants (67% and 46%, respectively) as illustrated in Figure 1.

**Figure 1:**
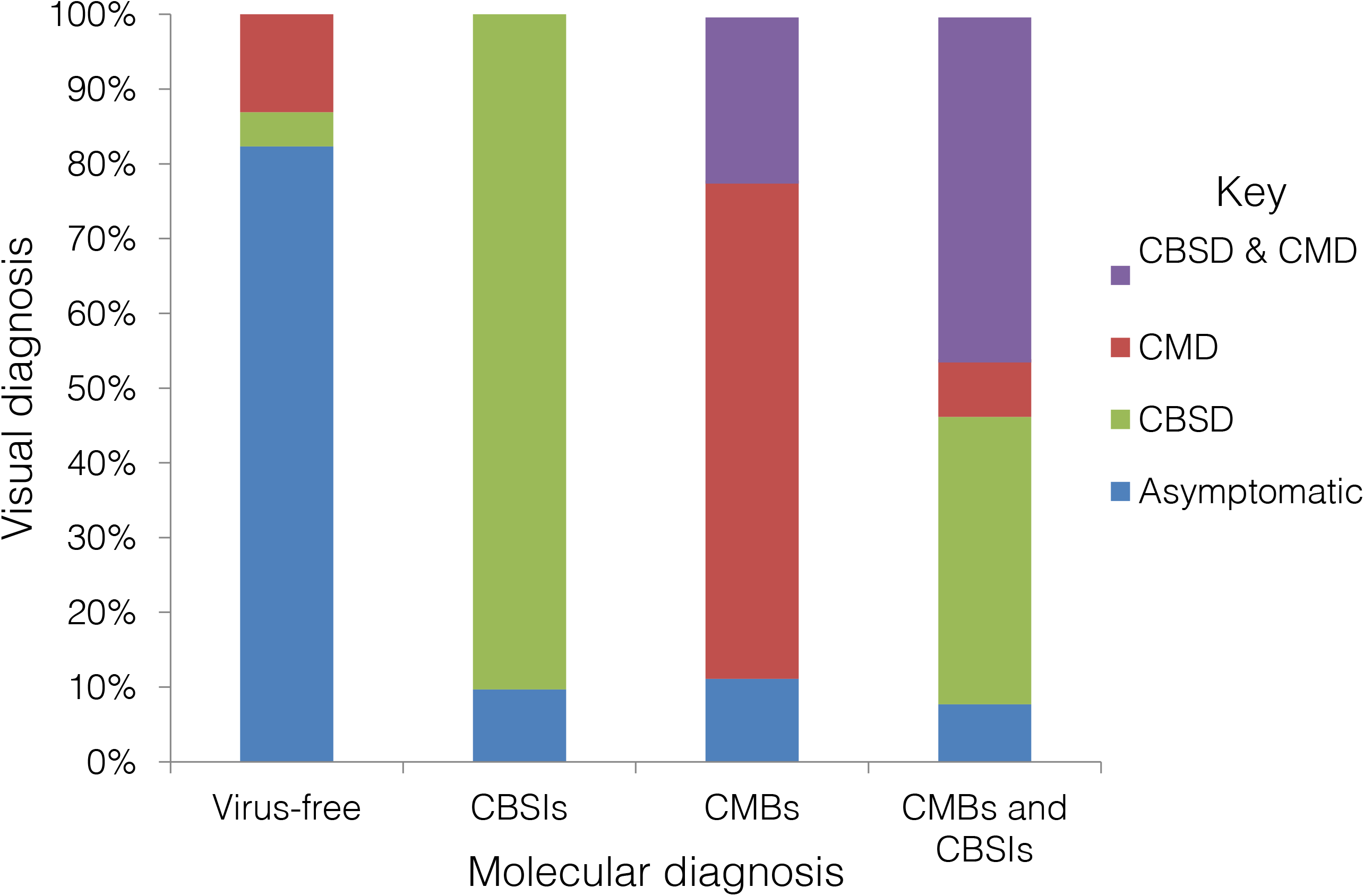
Comparison of visual and molecular diagnosis of CMD and CBSD on cassava plants.

Majority of the leaves that were co-infected with both CMBs and CBSIs were visually diagnosed to have symptoms of CBSD only (38%), suggesting that these plants could have latent CMD infection. Alternatively, the observed lower accuracy could also be due to presence of symptoms in other parts of the plants that were not sampled for virus confirmation. To confirm this, comparison of visual and molecular diagnosis was done for the leaves that were sampled for molecular diagnosis.

When the comparison of visual and molecular diagnosis was done based on the leaves that were sampled for molecular diagnosis the accuracy of the cassava experts to diagnose healthy asymptomatic plants increased slightly (from 82% to 91%) while accuracy for diagnosis of CBSD and co-infection decreased (from 90% to 64% for CBSD and from 46% to 13% for co-infection) and that for CMD was not affected, (Figure 2).

**Figure 2:**
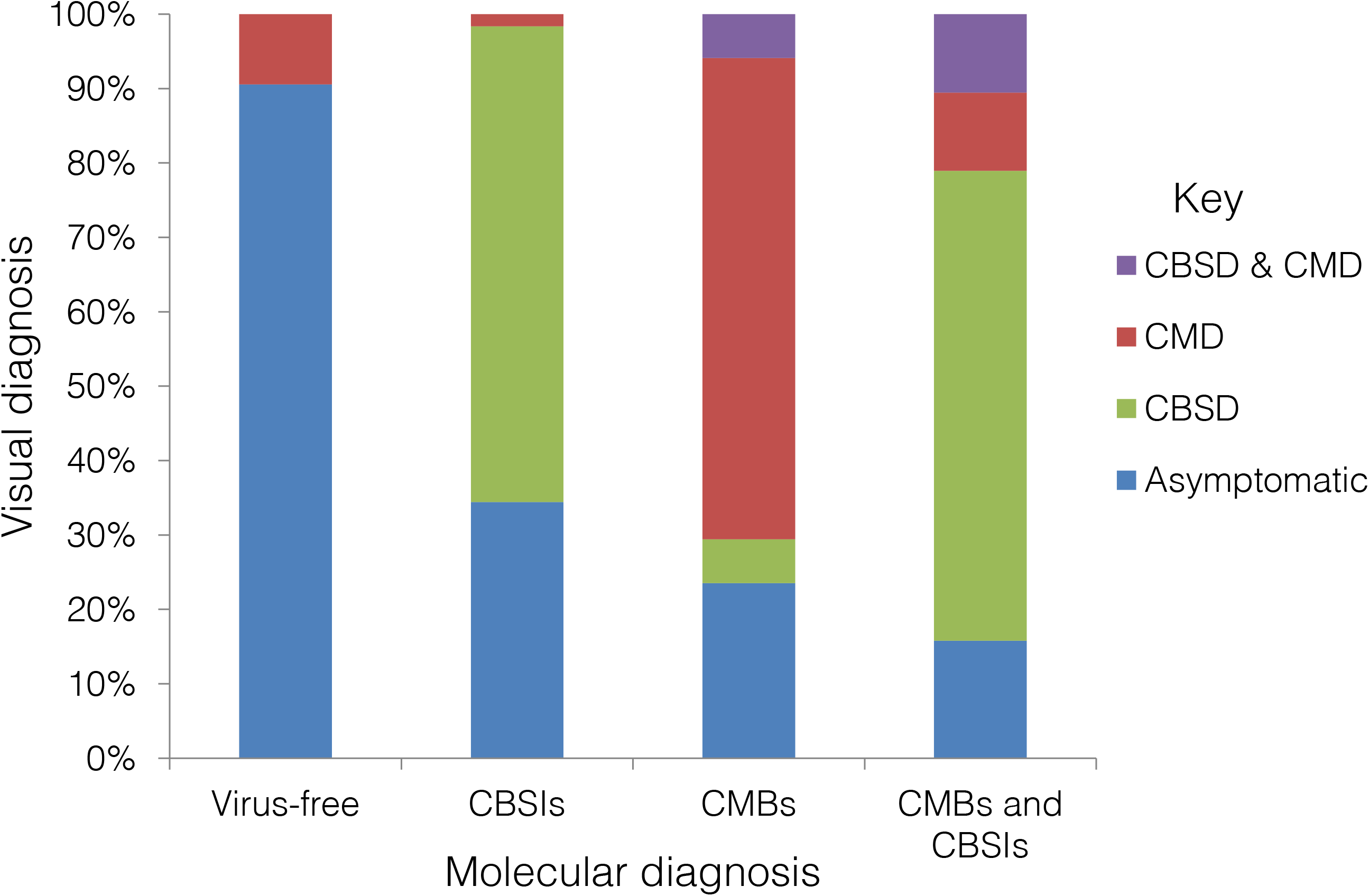
Comparison of visual and molecular diagnosis of CMD and CBSD on cassava leaves.

For the leaves that were infected with CMBs or CBSIs, most of the misdiagnosis were due to leaves that appeared to be asymptomatic, 34% for CBSD and 24% for CMD infected leaves. This was thought to be due to latent infection that has not resulted in disease symptom expression as previously suggested. However, the majority of leaves that were co-infected with viruses causing CMD and CBSD were visually diagnosed to have symptoms of CBSD only (75%), suggesting that the disease symptoms could mask each other and hence be difficult to distinguish. It can thus be concluded that the cassava experts could diagnose the presence of symptoms for CMD and CBSD with a high degree of accuracy in cassava plants containing viruses causing either CMD or CBSD but not in co-infected cassava plants. Increasing accuracy here would require the use of multiple leaves to inspect the whole plant.

### Comparison of the ability of Nuru and cassava experts in diagnosing symptoms of the viral diseases of cassava

The ability of *Nuru* to accurately identify symptoms of CMD, CBSD and CGM, and therefore diagnose presence of these diseases/conditions, was examined using 150 cassava leaves that were either asymptomatic or symptomatic for CMD, CBSD and CGM-damage. *Nuru* could identify asymptomatic leaves with a higher accuracy than symptomatic leaves (Table2).

**Table 2:** Accuracy of the cassava AI *Nuru* in diagnosing the presence of disease symptoms in cassava leaves.

| Condition | Number of plants | In-field assessment |  |
| --- | --- | --- | --- |
|  |  | No. of leaves accurately diagnosed | % of leaves accurately diagnosed |
| CMD | 90 | 53 | 58.90% |
| CBSD | 90 | 39 | 43.30% |
| CGM | 90 | 57 | 63.30% |
| Healthy | 30 | 26 | 86.70% |
| OVERALL | 300 | 175 | 58.30% |

The use of multiple leaves for diagnosis of the condition of a plant improved *Nuru’s* accuracy for diagnosing all the three conditions (CMD, CBSD and CGM) (Figure 3). The use of two leaves per plant significantly improved *Nuru’s* diagnosis capacity for CMD, CBSD and CGM as well as overall diagnosis of the whole plant. When the number of leaves was increased to three leaves per plant, there was a further slight increase in the symptom detection capacity of *Nuru* for CMD and CGM but there was a considerable increase in the diagnostic capacity for CBSD. For this reason, the use of multiple leaves for diagnosis of the condition of a plant was suggested as a means to improve the in-field capacity of *Nuru*.

**Figure 3:**
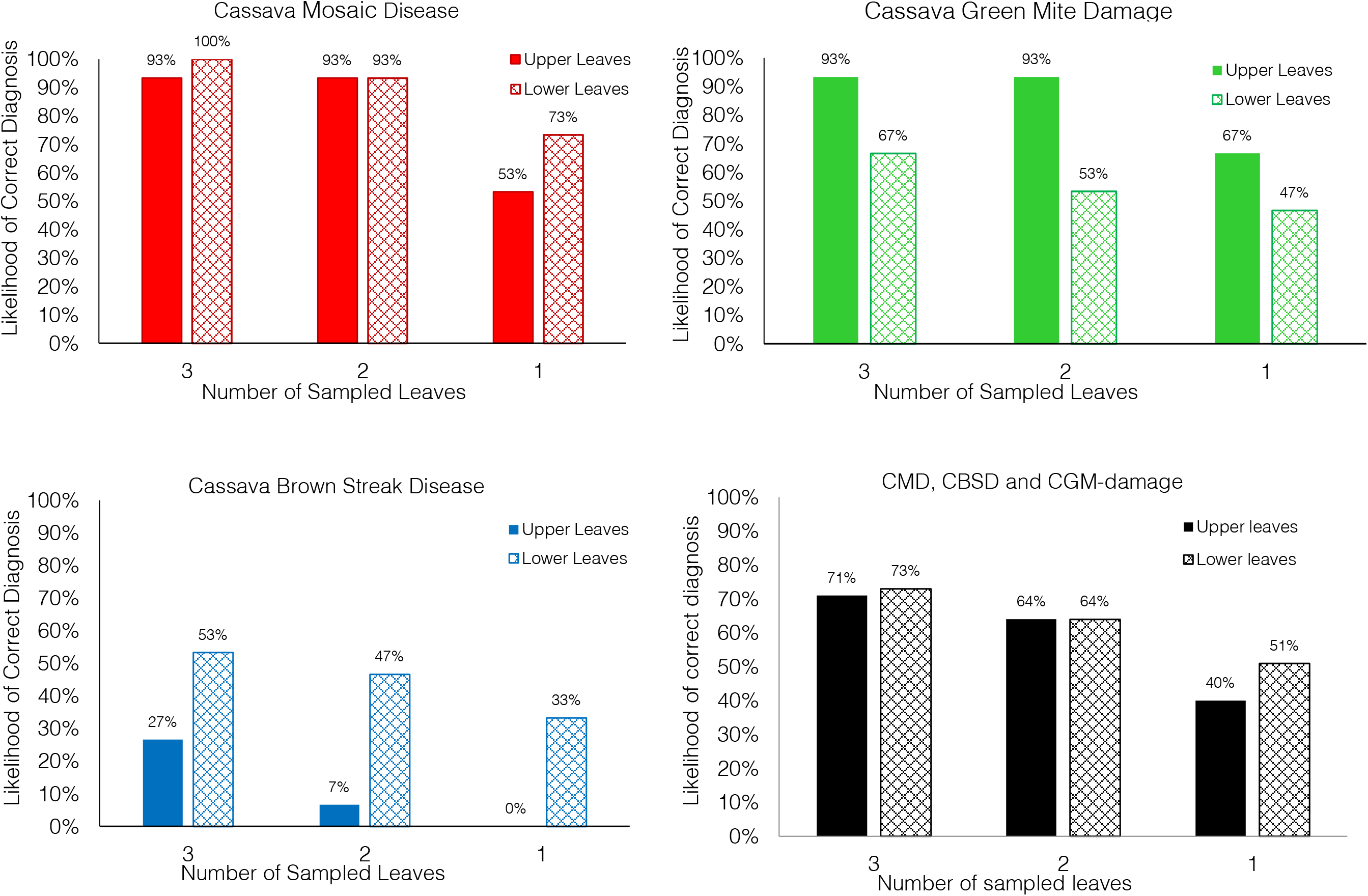
The likelihood of obtaining a correct diagnosis for CMD, CBSD and CGM - damage when using the cassava model in Nuru app for symptom detection based on the number of leaves sampled.

*Nuru’s* overall accuracy of identification of symptoms of the three conditions (CMD, CBSD and CGM) was not affected by the location of the sampled leaf. However, the location of leaves had an impact on *Nuru’s* accuracy for identifying symptoms of CGM and CBSD. Presence of CGM was more accurately identified using upper leaves while CBSD-symptoms were more accurately identified using lower leaves. The observed findings are not unusual since CGM are known to prefer the younger tender leaves which are usually the top leaves while CBSD symptoms are known to be clearer on the lower leaves (Nichols, 1950). When combined, the use of six leaves (3-upper and 3-lower) provided the highest likelihood of diagnosis (Figure 4) and this was the recommended number of leaves suggested for improving the accuracy of the cassava model in *Nuru*.

**Figure 4:**
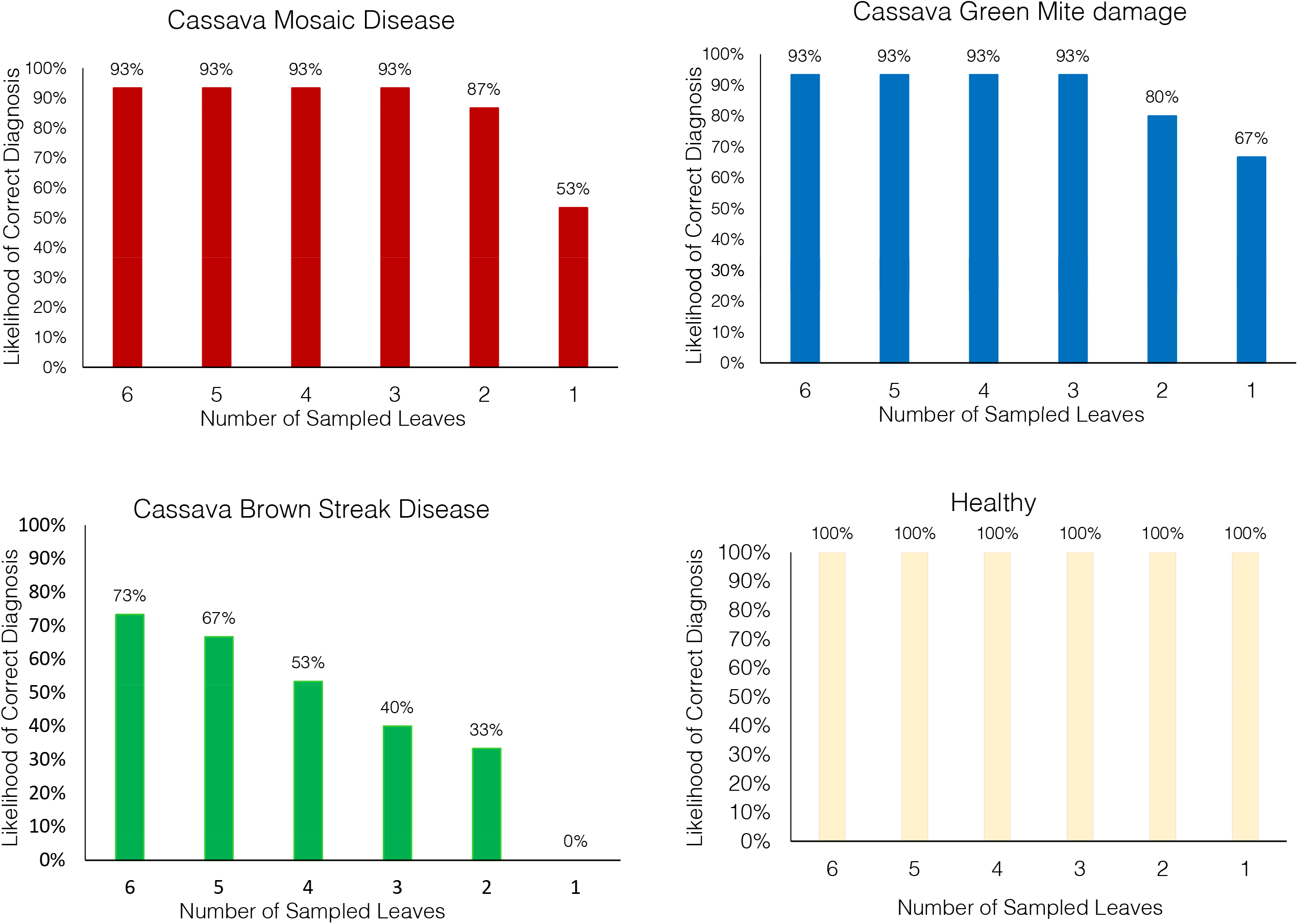
The overall likelihood of obtaining correct diagnoses for CMD, CBSD and CGM - damage when using the cassava model in Nuru app based on the number of leaves sampled.

### Comparison of the ability of Nuru, Agricultural extension agents and farmers to correctly diagnose symptoms of CMD, CBSD and CGM on cassava leaves

The cassava symptom recognition assessment tool was used to determine the ability of individuals and groups to accurately diagnose foliar symptoms. This was done by comparing the ability of researchers, agricultural extension officers and farmers (who have been previously trained on cassava pest and diseases and those who have not) to accurately identify symptoms of CMD, CBSD and CGM damage by using the cassava symptom recognition assessment tool. The researchers trained on cassava pests and diseases were able to identify symptoms of CMD, CBSD and CGM-damage with higher accuracy than untrained researchers, agricultural extension officers and farmers (Figure 5).

**Figure 5:**
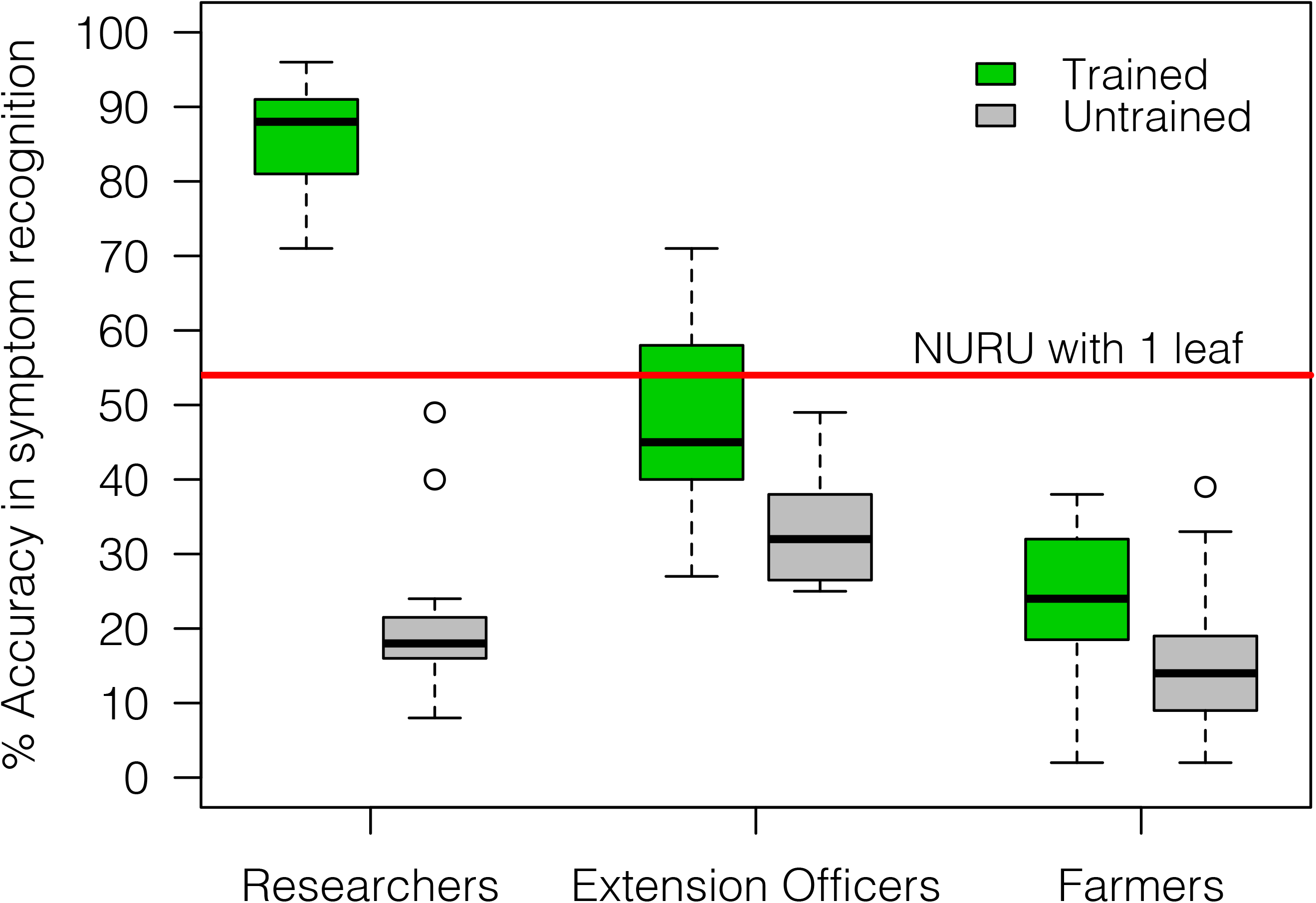
Comparison of accuracy in recognising symptoms of CMD, CBSD and CGM damage by the cassava AI Nuru, researchers, agricultural extension officers and farmers who have and have not previously received training on cassava diseases and pests.

The accuracy scores of symptom recognition by trained researchers were about four times higher than those of untrained researchers (Welch two-sample T-test p-value = 4.30E-11), and twice as high as those of trained agricultural extension agents and farmers (Tukey HSD p-value <0.005). There was a slight difference between the accuracy scores of trained and untrained agricultural extension officers, but both had higher accuracy scores than those of farmers. The observed difference could be due to the fact that the untrained agricultural extension agents and farmers were exposed to the symptoms of CMD, CBSD and CGM damage though their daily lives as most of them are involved in farming activities unlike the untrained researchers who are not involved in any activities related to cassava production. The large difference in accuracy scores of trained researchers in comparison to trained agricultural extension agents suggested that the vertical transfer of knowledge from researchers to extension agents is not very efficient, which is in part due to the lack of sufficient resources for training the very low frequency of training events involving both researchers and extension staff.

The symptom identification capacity of the cassava AI in *Nuru* was assessed using the cassava symptom recognition assessment tool and *Nuru* was able to identify disease symptoms with 54 % accuracy. The accuracy score of *Nuru* was similar to that of trained agricultural extension officers and slightly higher than that obtained from in-field diagnosis using one leaf as illustrated in Figure 3. As previously suggested, the use of six leaves was able to improve *Nuru’s* diagnosis accuracy (74 - 88 %) to a level similar to that of trained researchers (80 - 90 %), making *Nuru* as good as the experts. With six leaves, the diagnosis capacity of *Nuru* was significantly higher than that of trained agricultural extension agents (40 - 60%) and twice as high as that of trained farmers (20 - 32%).

### Evaluation of the teaching capability of Nuru-

The assessment tool was also used to evaluate the teaching capability of *Nuru* by determining the ability of agricultural extension officers and farmers trained on cassava pests and diseases to correctly identify symptoms of diseases after they used the *Nuru* app for about two weeks.

The accuracy scores of the agricultural extension officers were slightly higher than those of the farmers before the training and there was a significant increase in the ability of the extension agents to identify symptoms of CMD, CBSD and CGM after the training (Tukey HSD p-value <0.005) (Figure 6). However, this was not the case for the farmers where there was only a marginal change in their accuracy score after training, suggesting that training was not effective for the farmers. Further investigations are required to determine the most suitable method for training farmers and how these training can be conducted and their effect evaluated.

**Figure 6:**
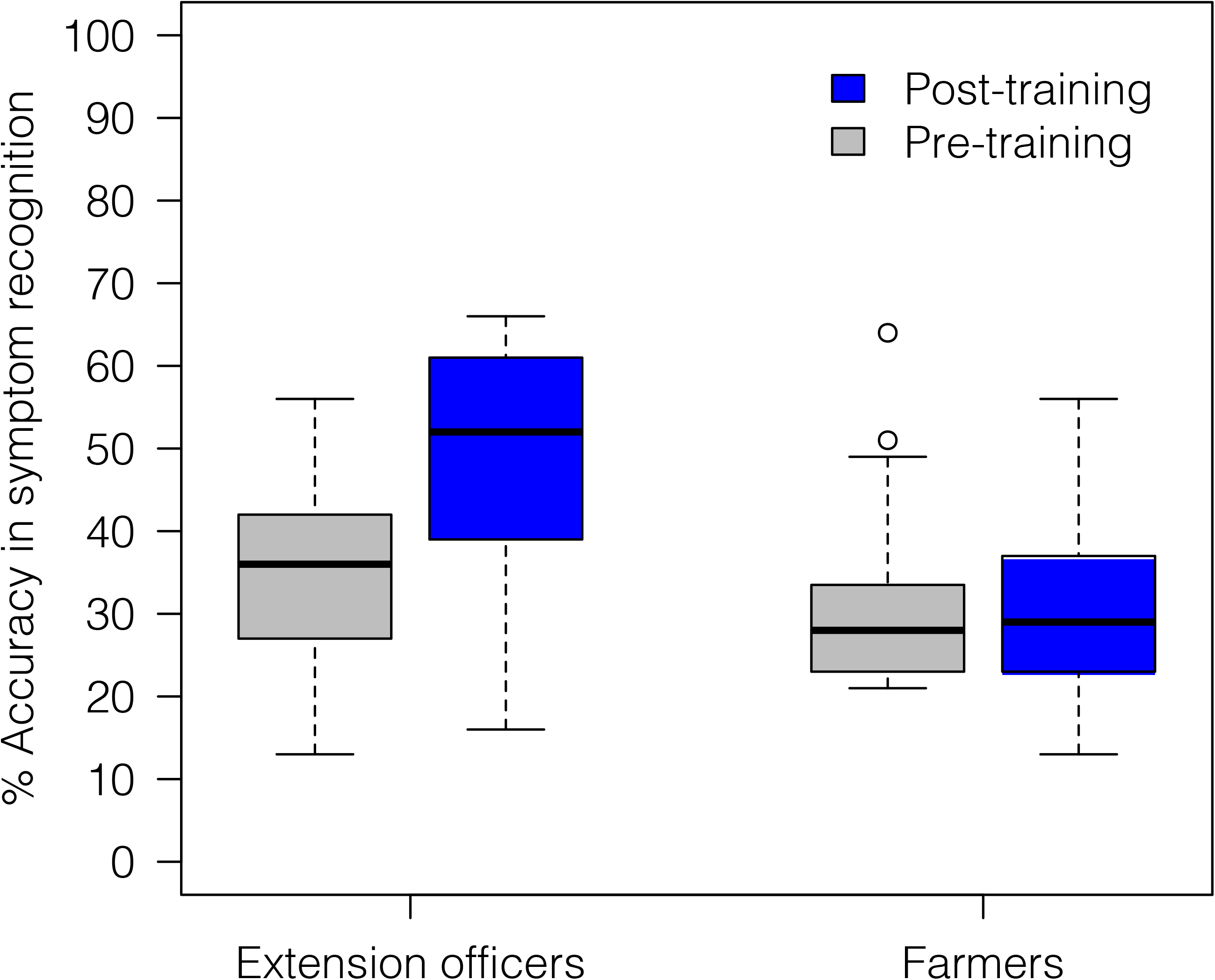
Comparison of the ability of agricultural extension officers and farmers to accurately recognise symptoms of CMD, CBSD and CGM-damage before and after being trained on cassava pests and diseases.

Even though there was no significant difference in the accuracy scores of the agricultural extension agents after being trained on cassava diseases and pests as well as using *Nuru*, the range of the accuracy scores (46-60%) was slightly narrower than that obtained from agricultural extension officers who had only received training without using *Nuru* (38-61%) (Figure 7).

**Figure 7:**
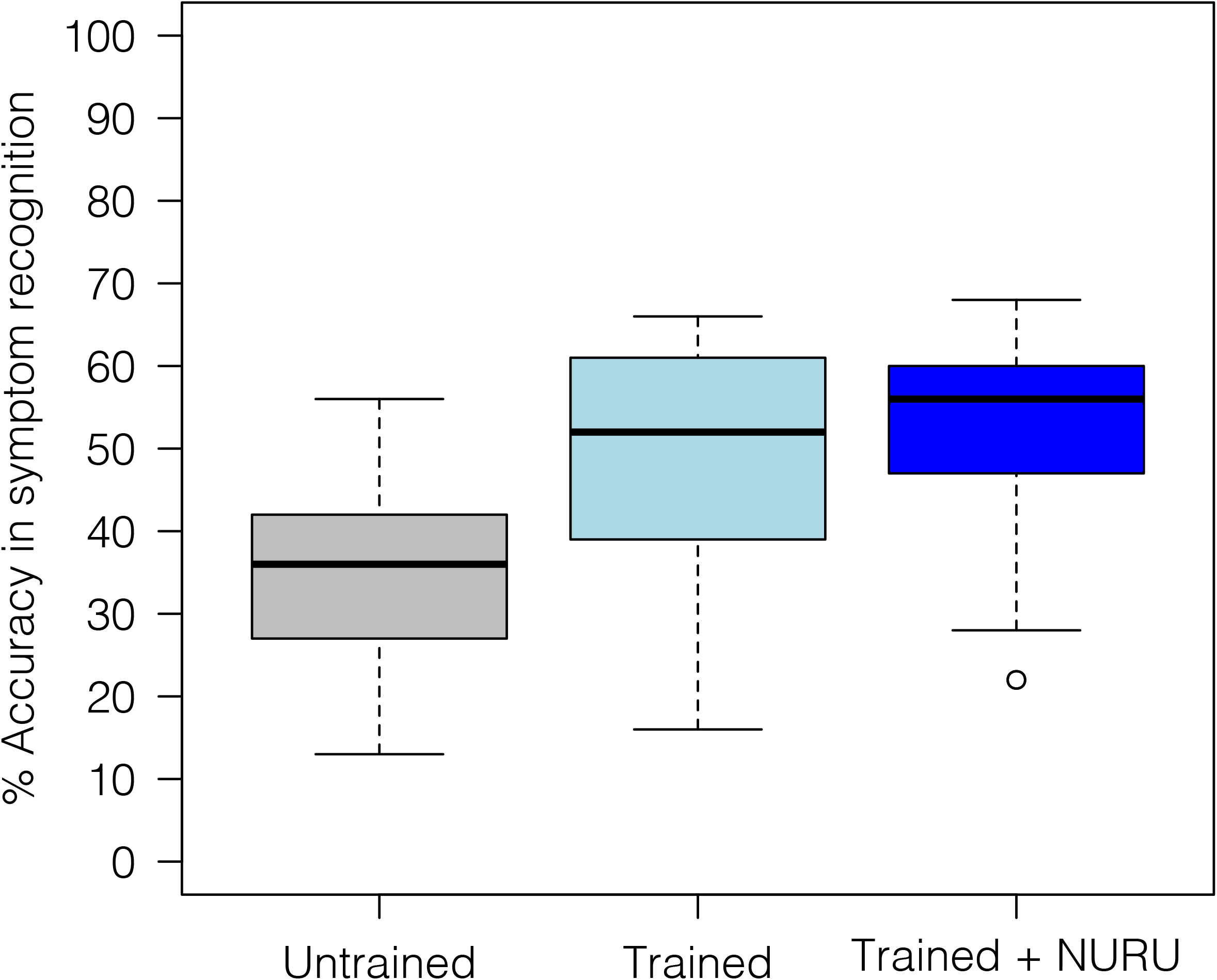
Comparison of the ability of agricultural extension officers to accurately recognise symptoms of CMD, CBSD and CGM-damage before and after receiving training on cassava pests and diseases as well as after using the cassava AI Nuru for diagnosing cassava diseases.

Overall, these results suggest that training delivers significant improvements in accuracy of disease diagnosis for farmers and extension officers, whilst the additional use of *Nuru* did not add to these improvements. Longer period of time might be required to observe changes in the capacity of agricultural extension agents to diagnose diseases themselves after using *Nuru*. Although the primary role of *Nuru i*s for rapid pest/disease diagnosis rather than training, it would nevertheless be worthwhile to investigate how its utility as a training tool might be improved.

## CONCLUSION

Development of artificial intelligence systems such as *Nuru*, which are based on machine learning models requires input from experts who are highly conversant with the features of the elements that are to be identified. Therefore, it is important to evaluate the expertise of these experts to ensure that the information used to train such models is as accurate as possible. Knowing the expertise of the ‘experts’ also provides a baseline with which the developed models can be evaluated.

The present work has shown that the cassava experts used to train and evaluate *Nuru* could identify the presence of CMD and CBSD symptoms with significantly higher accuracies than agricultural extension officers and farmers. The difference in symptom recognition capacity of the cassava experts, agricultural extension officers and farmers highlights the difficulty of diagnosing symptoms of CMD and CBSD, such that only well-trained individuals can do so effectively. Hence the ability of the cassava model in *Nuru* to have a similar capability for disease diagnosis as the cassava experts indicates that the model has been effectively trained and therefore has an important role to play as a tool for in-field diagnosis. The model that we tested here has already been improved and continuous improvements will be done as more data become available and so the diagnosis capacity of *Nuru* is expected to increase over time.

Even though the teaching capability of *Nuru* was not established, the app has the potential to be used as a training tool as it provides its users with opportunities to see the symptoms of the diseases present in the plant as it’s being used. The users can also view images with symptoms of identified diseases/types of pest damage from the library integrated into the app and they can also learn how to control and manage the pests and diseases identified from their plants through the advice given in the app. Current advice is formatted as simple blocks of text. This might be improved by developing more interactive training modules housed within the app. By enhancing the capabilities of *Nuru* to teach and train its users about the symptoms of the diseases affecting different crops one can provide a quick and cost-effective way of disseminating knowledge from researchers to agricultural extension agents and farmers. Furthermore, *Nuru* can provide an easily accessible means for ensuring continuous training of its users, thereby improving their skills in pest/disease identification and management.

On-going work in western Kenya indicates that the *Nuru* AI assistant is improving disease diagnosis skills and is having positive effects on communities of farmers who are using it. This suggests that efforts to scale out the use of *Nuru* across the cassava-growing regions of Africa will improve farmers’ recognition and knowledge of cassava diseases/pests, which will contribute to improved disease/pest control ultimately leading to increased yields and strengthened food security.

## ACKNOWLEDGMENTS

This work was financed through an ‘Inspire Challenge’ grant from the BigData Platform of the Consultative Group on International Agricultural Research (CGIAR) and the Huck Institutes of the Life Sciences at Penn State University. The contributions of J.L. were supported through the Roots, Tubers and Bananas (RTB) research programme of the CGIAR. The support of the Ministries of Agriculture in Tanzania and Kenya is recognized, as are the contributions of farmers and extension staff in Mkuranga District, Tanzania and Busia County, Kenya.

## AUTHOR CONTRIBUTIONS

L.M, N.M, M.N, A.R, A.K, H.M and P.M collected the data. J.L and D.H supervised the project.

L.M prepared the manuscript with support from J.L and D.H.

