## Supplementary A for "Evaluation of the accuracy of a smartphone-based artificial intelligence system, PlantVillage *Nuru,* in diagnosing of the viral diseases of cassava"

***Molecular diagnosis of CMD and CBSD***

<u>Extraction of nucleic acid from investigated leaves</u>

Prior the PCR amplification the cassava leaves were dried, ground and total nucleic acid was extracted from the ground leaves using the cetyl trimethyl ammonium bromide (CTAB) extraction procedure adapted from Maruthi (2002).

Briefly, 1 mL of the CTAB buffer (containing 2.0%, w/v CTAB, 2.0% PVP, 25 mM EDTA, 2.0 M NaCl, 100 mM Tris HCL pH 8.0 and 0.2% β-mercaptoethanol which was added immediately before extraction) was added into the dried cassava leaves. The leaves were then ground and incubated at 65°C for 15 minutes to lyse the cells and facilitate the separation of polysaccharides and polyphenols from the cellular material. An equal amount of chloroform:isoamyl alcohol (24:1) was added to the cell lysate to separate the nucleic acids from the cell lysate. The nucleic acid was then precipitated from the solution by adding 0.6x volume of cold isopropanol and the samples were incubated at -20°C for 30 minutes prior to centrifugation at 13,000 rpm and 4°C for 10 minutes to pellet the nucleic acids. The supernatant was discarded and the nucleic acid pellet was washed twice by adding 700 µL of 70% ethanol, vortexing and incubating at -20°C prior to centrifugation at 13,000 rpm for 5 minutes. Afterwards, the ethanol solution was removed, the nucleic acid pellet was air-dried and then resuspended in 100 µL Tris-EDTA buffer (1x) . The quantity and quality of the nucleic acid extracts were determined by spectrometry at 260 nm.

PCR amplification of viruses that cause CMD

The nucleic acid virus that cause CMD in the coastal region (EACMBs) was amplified using the procedure and primers described by Ndungure et al. (2005). Briefly, about 20 ng of the total nucleic acid was added into he PCR master mix containing One Taq 2x Master mix with standard buffer (M0482S, New England Biolabs) and 200 nM of each forward and reverse primers. The PCR amplification was done using Veriti thermocycler (Applied Biosystems) at the following cycling conditions: initial DNA denaturation at 94°C for 2 min, followed by 30 cycles of denaturation at 94°C for 30 seconds, annealing at 55°C for 30 seconds and extension at 68°C 40 seconds followed by a final extension at 68°C for 10 minutes. The PCR products were analyzed by agarose gel electrophoresis using 1% (w/v) agarose gels and 1X TAE buffer. The DNA products were stained by GelRed nuclec acid stain (Biotium, California, USA) and the gels were viewed and photographed using the Syngene GBox system (Syngene, Cambridge, UK). Samples containing DNA bands of about 550 bp were considered as CMV positive results.

PCR amplification of viruses that cause CBSD

Detection of the virus that cause CBSD, CBSVs and UCBSVs, was done by real-time RT-PCR using Taq chemistry using the method and primers described by Shirima et al. (2017). Briefly, 4 µL of the template nucleic acid was added into PCR reaction mixtures containing 1x PCR buffer, 5.5 mM $MgCl_2$, 0.5 mM dNTPs, 300 nM primer, 100 nM probe, 30 nM Rox reference dye, 0.625 Units of Taq DNA polymerase and 0.4 Units of M-MLV- reverse transcriptase into a 25 µL reaction. The Taq DNA polymerase and reverse transcriptase were obtained from Life Technologies (California, USA) while all the reagents in the PCR master mix were obtained from IDT (California, USA). An internal control (Cytochrome oxidase 1), a negative control (no-template) and positive controls (samples which have been previously tested to have the virus of interest) were added.

The amplification reactions were done using Stratagene MX3000P (Agilent Technologies, New Jersey, USA) with the following thermo-cycling conditions: 30 minutes incubation at 48˚ for reverse transcription, initial denaturation of the cDNA at 95˚C for 10 minutes, 40 cycles of denaturation at 95˚C for 15 seconds and annealing and extension step at 60˚C for 1 minute. Fluorescent data were collected during the 60˚C step using Stratagene MxPro Real-time QPCR software version 4 (Agilent Technologies, New Jersey, USA). Based on the amplification curves, samples with cycle threshold (Ct) values below 36 were considered as positive results.

<u>References</u>

Maruthi, N., Colvin, J., Seal, S., Gibson, G., and Cooper, J. 2002. Co-adaptation between cassava mosaic geminiviruses and their local vector populations. *Virus Research* 86:71-85.

Adams, P., Abidrabo, P., Miano, W., Alicai, T., Kinyua M, Clarke, J., Macarthur, R., et al. 2012. High throughput real-time RT-PCR assays for specific detection of cassava brown streak disease causal viruses and their application to testing planting material. *Plant Pathology*. 62: 233–242.
