## Supplementary B for "Evaluation of the accuracy of a smartphone-based artificial intelligence system, PlantVillage *Nuru,* in diagnosing of the viral diseases of cassava"

***Development of the cassava symptom recognition assessment tool -***

The cassava symptom recognition assessment tool as scoring system for 170 images of cassava leaves which are asymptomatic and symptomatic for CMD and CBSD as well as CGM-damage. The system was developed by 10 cassava experts who have been working on cassava pests and diseases for 2 to over 10 years. The experts individually scored 170 images of cassava leaves based on the symptoms they recognized on the leaves and the scoring key illustrated in Table 1.

The scores for each image were compared to identify images that were diagnosed differently by more than three individuals. These images were discussed to determine the disease symptom that was present on the leaves and the agreed symptom, together with symptoms of the images that all the individuals in the group agreed on, were considered an the consensus symptom.

The consensus symptoms were converted into a numbers representing a numerical code for each of the observed condition as illustrated in Table 1.

The numerical codes were used to score the condition of the 170 images in an excel template which allowed for comparison of the codes entered by the assessed individuals to those representing the consensus symptoms identified by the experts. The comparison was based on counting the number of times an individual scored an image similarly to the expert and using this count to calculate the percentage accuracy for diagnosing symptoms of cassava diseases. The excel template also enabled identification of images that were incorrectly diagnosed, based on the expert diagnosis, and conditions which were misdiagnosed.

**Table 1.1: Numerical codes to identify the conditions of the leaf images used for the development of the cassava symptom recognition assessment tool.**

| Condition | Abbreviation | Score |
|---|---|---|
| Cassava mosaic disease | CMD | 1 |
| Cassava brown streak disease | CBSD | 2 |
| Cassava green mites | Mites | 3 |
| Brown leaf spots | BLS | 4 |
| Fungal-like infection | FLI | 5 |
| Healthy | - | 6 |
| Other | - | 7 |
| Not sure | - | 8 |
